# How collectively integrated are ecological communities?

**DOI:** 10.1101/2022.12.29.522189

**Authors:** Yuval R. Zelnik, Nuria Galiana, Matthieu Barbier, Michel Loreau, Eric Galbraith, Jean-François Arnoldi

## Abstract

Beyond abiotic conditions, do population dynamics mostly depend on the species’ direct predators, preys and conspecifics? Or can indirect feedbacks that ripple across the whole community be equally important? Here we show that the spectral radius of a community’s interaction matrix controls the length of indirect interaction pathways that actually contribute to community-level dynamical patterns, such as the depth of a perturbation’s reach, or the contribution of biotic processes to realized species niches. The spectral radius is a measure of collectivity that integrates existing approaches to complexity, interaction structure and indirect interactions, while also being accessible from imperfect knowledge of biotic interactions. Our work provides an original perspective on the question of to what degree communities are more than loose collections of species or simple interaction motifs; and explains when reductionist approaches focusing on particular species and small interaction motifs, ought to suffice or fail when applied to ecological communities.

## Introduction

Ecological communities comprise vast networks of interacting species, that greatly vary in their richness and connectivity (Pimm, 1984; Montoya *et al*., 2006; Agrawal, 2001; Brown *et al*., 2001). To understand and predict their behaviour it is tempting to take a reductionist approach, breaking-down complex communities into small parts (predator-prey pairs, competitors within a same niche, etc.). For instance, we might hope that, to understand a population’s dynamical response to environmental change, one should look at this species response traits (Lavorel & Garnier, 2002), then those of the other species with which it interacts directly and most strongly. Perhaps, but to a lesser extent, one would also consider indirectly connected species. Here one would rely on the validity of incremental causality, as we gradually build-up a chain of causal links between various interacting units. But a radically different, holistic, perspective would be to view a species’ response to environmental change as a collective response of the whole ecosystem in which it is embedded (Patten, 1982).

Explicitly or implicitly, for decades ecologists have argued whether reductionist or holistic perspectives are most appropriate (Loreau, 2020). This debate is often traced back to the opposition, regarding plant communities, between the holistic view of Clements, and the parsimonious individualistic perspective of Gleason (Lefkaditou, 2012). Clements argued that plant associations should be understood as high-level biological entities, comparable to actual organisms, so that species are best understood through their functions within a whole (Clements, 1916). Gleason claimed that plant communities are mere collections of individual species and gave little importance to the interactions between them (Gleason, 1926). This dichotomy has carried on, with notable ideas such as Lovelock’s Gaia theory proposing that the biosphere should be viewed as a super-organism (Lovelock & Margulis, 1974), a perspective that profoundly contrasts with the ideas behind the use of Species Distribution Models, that aim to predict species ranges with few key environmental variables (Soberón, 2007).

But there also exists a middle way that embraces at once some of the above reductionist and holistic views of ecological communities (Lefkaditou, 2012). We can model them as high-dimensional dynamical systems, where variables represent species abundances whose dynamics are coupled by interaction terms. Those interactions are encoded in a matrix, so that ecological communities are mapped to a rich class of mathematical objects whose properties can reveal emergent features of the system –that in turn influence the behaviour of its parts (Levins & Lewontin, 1982). Robert May famously showed, using random matrix theory, that stability was virtually impossible past a complexity threshold (May, 1972). This thought-provoking result, contradicting heuristic ideas of their time, opened up a fruitful line of research, looking for interaction structures allowing complex communities to persist (Allesina & Tang, 2015), thus asking questions about ecological structure and dynamics in terms of matrix features (Novak *et al*., 2016). In particular, the eigenvalues of an interaction matrix (its spectrum), reveal the dominant modes by which biotic iterations influence population dynamics (Trefethen & Embree, 2020). Eigenvalues are used to determine local stability (Allesina & Tang, 2012; Neubert & Caswell, 1997; Tang & Allesina, 2014), the spectral radius of adjacency matrices reflects the nestedness of bipartite networks (Staniczenko *et al*., 2013), and from the singular value decomposition of interaction matrices, the likelihood of species coexistence can be derived (Rohr *et al*., 2014; Grilli *et al*., 2017). Here, using these mathematical objects and techniques, we address the reductionist/holistic opposition, and show that it can be considered a continuous axis (Liautaud *et al*., 2019), along which different communities position themselves depending on their complexity and structure (Allesina & Pascual, 2008). We will explain what this positioning tells us about their observable behaviour (e.g. response to perturbations), and discuss how this could be assessed empirically. To do so, we revisit a classic notion of community ecology, that may seem unrelated at first: indirect interactions between species (Bender *et al*., 1984).

The existence of indirect interactions between species indeed challenges the reductionist/individualistic approach to ecological communities (Menge, 1995; Abrams *et al*., 1996). These interactions are mediated via one or several intermediate populations (Wootton, 1994) and form long, and numerous, pathways across the community (Puccia & Levins, 2013), generating a multitude of confounding causal pathways between its constituent species. One may think of the feeding chains that couple fungi, bacteria and invertebrates in soil food-webs (Neutel *et al*., 2002), or the indirect interactions between fish and plants via dragon-flies whose larvae are eaten by fish and whose adults prey on plant pollinators (Knight *et al*., 2005). Indirect interactions can couple biomes, determine the loops that control the stability of food-webs, and impair our capacity to predict a species responses to a given perturbation (Yodzis, 1988). Indirect interactions thus generate intricate interconnections, leading to emergent community behavior that is different from a collection of populations or isolated interaction motifs (Loreau, 2020).

Here we relate the importance of indirect interaction pathways to the spectral radius of interaction matrices (their largest eigenvalue modulus). Taking a dynamical system’s perspective, we compare direct interactions between species pairs, to their long-term –net– interactions (Montoya *et al*., 2009; Novak *et al*., 2016); to then explain how the latter integrates all indirect interaction pathways between them. We demonstrate that the spectral radius of the interaction matrix (once properly normalized by self-regulating forces) determines the length of indirect pathways that contribute to net-interactions and thus to long term-community dynamics and patterns. We call this length “interaction horizon”, a notion which mirrors the “environ” concept proposed by Patten (1982) from the era of theoretical ecosystem ecology. As we will see, when interaction horizon diverges, this does not imply unstable behaviour, but the break-down of incremental causal thinking.

On simulated communities we will illustrate that the spectral radius of an interaction matrix explains the occurrence of intuitive signatures of collective community behavior, such as the depth of a perturbation’s reach, the degree of temporal unpredictability of a community’s response to environmental change, or the contribution of biotic processes to realized species niches. We therefore propose the spectral radius of the interaction matrix as a measure of the collectivity of a community. From spectral radius bounds, in which May’s complexity measure plays a central role, we show how to quantify the dynamical role of interaction structures. For instance, in cascade food-webs models (Pimm *et al*., 1991), we show that trophic efficiency increases collectivity, and so does the nestedness of bi-partite (e.g. plant-pollinator) networks (Staniczenko *et al*., 2013). Our measure of collectivity thus embraces the somewhat disconnected existing approaches to community complexity, organizational integration and indirect interactions.

Our work clarifies when pragmatic reductionist perspectives, focusing on particular species and small interaction motifs can, at least in principle, reliably scale-up to the community-level, or on the contrary, when there are fundamental obstacles facing such approaches (Bergelson *et al*., 2021; Orr *et al*., 2021).

### Collectivity and the interaction horizon

Here we provide an introduction to a collectivity parameter focusing on intuitions, while in Box 1 we give a more formal and general derivation. We then apply those ideas on a simple resource-consumer model.

Start from a community’s interaction matrix *A* = (*A*_*ij*_), with *A*_*ii*_≡0. *A*_*ij*_ is a non-dimensional number that quantifies the direct interaction of species *j* on species *i*. It is crucial to see *A*_*ij*_ as a relative interaction strength: the ratio of inter-over intra-specific interactions. We emphasise this seemingly technical detail because it is key to properly define a notion of indirect interactions. If interactions had units, indirect interactions of different orders would themselves have different units, making them incomparable. Studying interaction matrices that have a constant diagonal is common place in theoretical ecology, yet it is rarely actually justified (May, 1972; Allesina & Tang, 2012; Jacquet *et al*., 2016). This property arises naturally when considering relative interaction strength, which for us is a fundamental prerequisite (see Box 1 and example of eq. 6 below).

Following (Puccia & Levins, 1991; Neutel *et al*., 2002), the indirect interaction of second order between species *j* and *i* through a third species *k* is *A*_*ik*_ *×A*_*kj*_, the product of the direct interaction of species *j* on species *k* with the direct interaction of species *k* on species *i*. More generally, an interaction pathway of length *n* between species *i* and *j* is 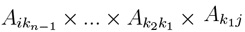, where the intermediate species *k*_1_, …, *k*_*n*_ need not all be different (loops are possible). Importantly, the sum over all such interaction pathways coincides with the element of *A*^*n*^:

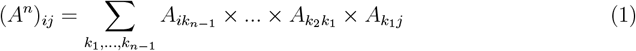

If all the numbers *A*_*ij*_ are strictly smaller than one, the magnitude of indirect interactions will decay exponentially as *n* grows. Conversely, if the interaction network is sufficiently connected, the number of interaction pathways between species *i* and *j* (the number of terms in the sum) will also increase exponentially. It is therefore not clear if the sum of all indirect interactions will necessarily vanish as their order grows, even if direct interactions are individually weak.

To see why Eq. (1) will appear in community models, consider the steady state condition of a Lotka-Volterra system (see Box 1 for a general case):

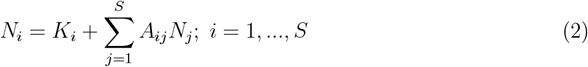

Here *K*_*i*_ –the carrying capacity of species *i*– has units of biomass and encodes the environmental conditions perceived by that species on its own. If we introduce the vectors *K* = (*K*_*i*_) and *N* = (*N*_*i*_), Eq. (2) can be written in compact form as

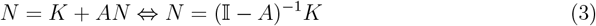

Thus the species’ intraspecific features *K*_*i*_ intertwine via the matrix (𝕀*−A*)^*−*1^ to determine the actual species abundances in the community context. For instance, a favourable environment (a large *K*_*i*_) will not imply a large abundance if the environment is also favourable to a competitor. The matrix (𝕀*−A*)^*−*1^ encodes all such effects, that is, all *net interactions* between species. If we had instead repeatedly applied Eq. (2) on itself we would have written a series highlighting the contribution of indirect interactions pathways, as defined in Eq. (1):

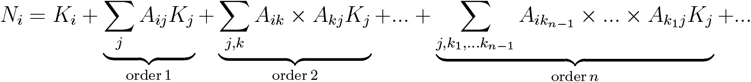

Since this last expression should be equivalent to Eq. (3), and this for all vectors *K*, we arrive at a classic matrix identity, known as Neumann’s series (Reed *et al*., 1972),

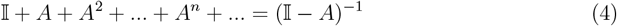

This series converges only under some specific conditions. When it does not converge, this means that we cannot meaningfully decompose net interaction as a sum of indirect interaction pathways.

The criteria for convergence gives us both a measure of the importance of indirect interactions, and a *definition* of collective integration. To derive this criteria, we first need to measure the magnitude of the various terms of the series, representing the overall strength of indirect interactions of all orders. This amounts to defining a matrix norm for each term ||*A*^*n*^||, and see how this norm changes with the order *n*. Consider

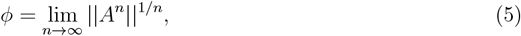

that is, the rate of growth of the norm ||*A*^*n*^|| as *n* grows. If *ϕ <* 1, as *n* grows, the overall contribution of indirect pathways will *eventually* decrease exponentially as *ϕ*^*n*^. If *ϕ >* 1, the sum over arbitrarily long pathways can be arbitrarily large (cf. Fig. 1).

**Figure 1.**
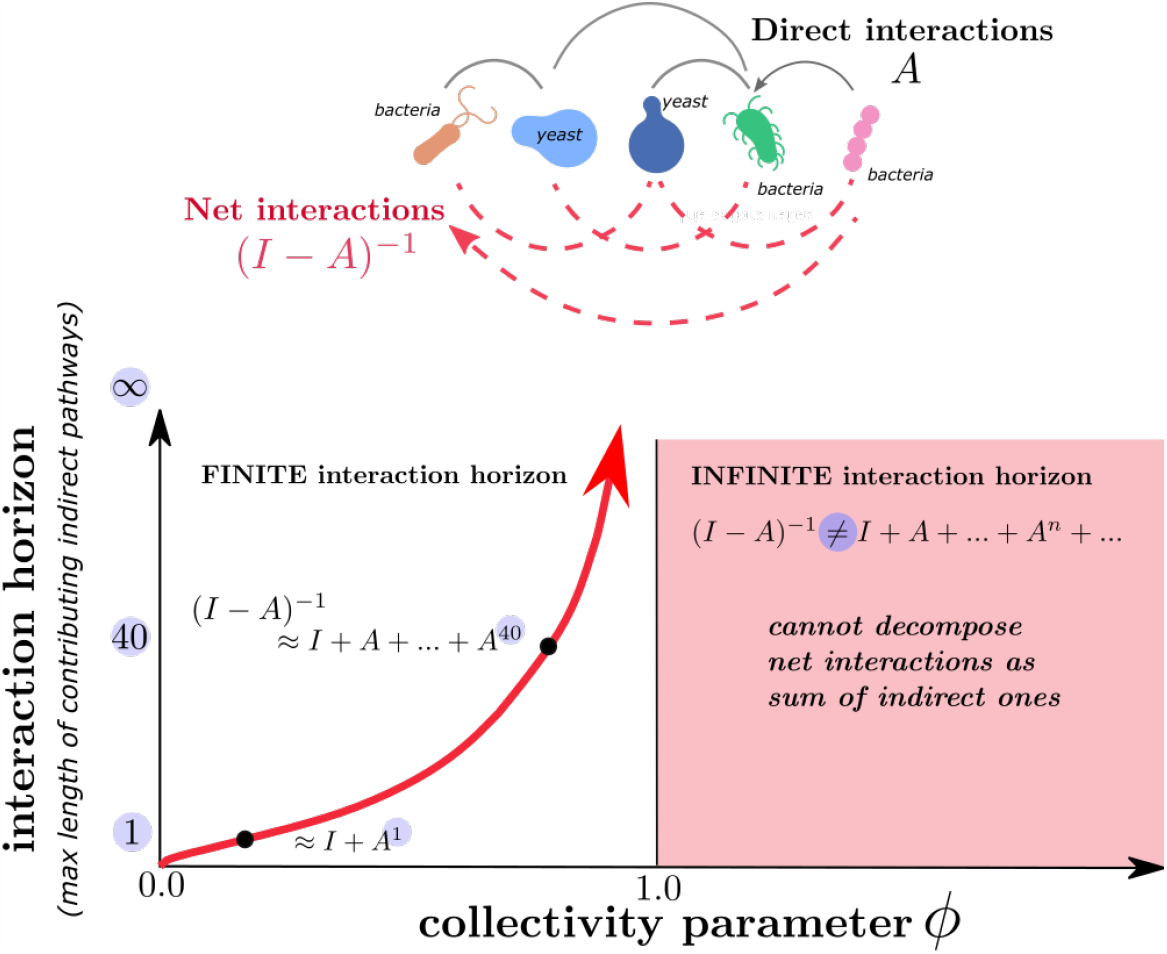
The interaction horizon is the maximal length of indirect interactions pathways that substantially contribute to net interactions (illustrated here in a hypothetical yeast-bacteria community). The horizon is directly determined by the collective parameter as log logϵ/*logϕ*, where *ϵ* < 1 is an arbitrary threshold value. The horizon gives the lowest order of interactions for which the maximal contribution is negligible (that is, smaller than *ϵ*), and it diverges as *ϕ* approaches 1. Beyond this point it no longer makes sense to decompose net interactions as a sum of indirect pathways.

Remarkably, *ϕ* does not depend on the particular choice of matrix norm. It is an intrinsic feature of the interaction matrix *A*: its spectral radius, the largest absolute value of its eigenvalues (Trefethen & Embree, 2020).

Here we propose an ecological interpretation of the spectral radius *ϕ* of a given interaction matrix. We call *ϕ* the *collectivity parameter*, because it determines the *interaction horizon* of species: the maximal length of interaction pathways that contribute to their net interactions (see Fig. 1). For systems for which *ϕ >* 1, the interaction horizon is infinite, signaling the breakdown of the reductionist method of decomposing net effects into indirect interaction pathways, which we see as a reflection of the highly collective integration of such communities.

**Example:** Let’s see how the above ideas play out in a simple resource-consumer model (see also Appendix S2). Let *N*_*C*_ and *N*_*R*_, be the abundance of consumer and resource, respectively, which are assumed to follow, as in (Galiana *et al*., 2021):

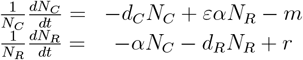

Here *d*_*C,R*_ represent intra-specific interactions, *α* is the attack rate, *ε* is trophic efficiency while *m* and *r* are intrinsic mortality and growth rate of both species. Following the general derivation of Box. 1, the non-dimensional interaction matrix is

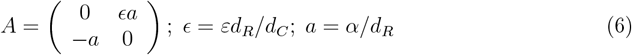

This matrix has purely imaginary eigenvalues 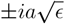, so that the collectivity parameter is 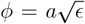, proportional to the attack rate. Because eigenvalues are imaginary, stable steady states exist^1^ for any values of *ϕ*. The inverse matrix –that determines net interactions– can be written in terms of *ϕ*:

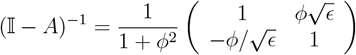

The net interaction between resource and consumer is thus proportional to *ϕ/*(1 + *ϕ*^2^). This non monotonous function of *ϕ* (and thus of attack rate) increases until *ϕ* = 1, before decreasing. We deduce that at high collectivity, net and direct interactions become anti-correlated. This counter-intuitive phenomenon cannot be understood when considering only a few, albeit long, indirect interaction pathways, because such a decomposition would only converge in the phase where net and direct interactions go hand in hand.

### Three signatures of collective integration

We now introduce three signatures of collective integration, which could conceivably be observed empirically. Not all three would be indicative of collectivity in a given system, but taken together they apply to a broad spectrum of ecological scenarios.

We look for those signatures on a set of Generalized Lotka-Volterra (GLV) model communities, taken in a steady state following community assembly from a random species pool.

In simulations we consider a gradient *y* of interaction strength (and heterogeneity), with 50 different values of overall interaction strength between 0.02 and 1, each with 100 random communities, making 5000 communities in total. Each starts from a pool of *S* = 50 species, and we set 80% of interactions to zero to have sparse interactions networks. To parameterise interaction strength, we follow (May, 1972; Allesina & Tang, 2012; Bunin, 2017; Barbier *et al*., 2018), and define three parameters of random interactions: 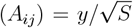 and corr(*A*_*ij*_, *A*_*ji*_) =*−*1. This leads to anti-correlated interactions between species, that are increasingly negative and varied. In this way we could generate communities with a collectivity parameter *ϕ* ranging between 0 and ∽2, which we use to showcase generic aspects of low and high collective integration in species-rich communities.

Details about simulations, and an additional signature (that we call “growth of effective connectance”) are given in Appendices S5 and S6.

### Perturbation depth

Collective integration means that species are interdependent, so that a perturbation targeted on a given species will likely propagate deep into the community (Bender *et al*., 1984). Experimentally one could remove a species, and monitor the responses of others, as a function of their interaction distance *d*(*i, j*) from the removed species (*d*(*i, j*) = 1 if *j* interacts with *i, d*(*i, j*) = 2 if *i* and *j* are indirectly connected via a third species, etc.). Denoting *N*_*j\i*_ the long term abundance of species *j* after removal of species *i*, one can quantify *perturbation depth* as

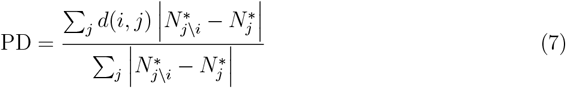

which we can average over all species removal experiments in that community.

In Fig. 2a we demonstrate a good agreement between this observable signature of collective integration and the collectivity parameter *ϕ*. As collectivity grows, the brunt of the perturbation effect is shared with more distant species, and not only supported by those directly in contact with the removed node (Fig. 2b). An obvious caveat of perturbation depth is that it only applies to sufficiently sparse networks – if all species are connected, this notion is not well defined. We show in Appendix S6 how this limitation can be overcome, by defining a distance function *d*(*i, j*) that is based on the quantitative values of interaction.

**Figure 2.**
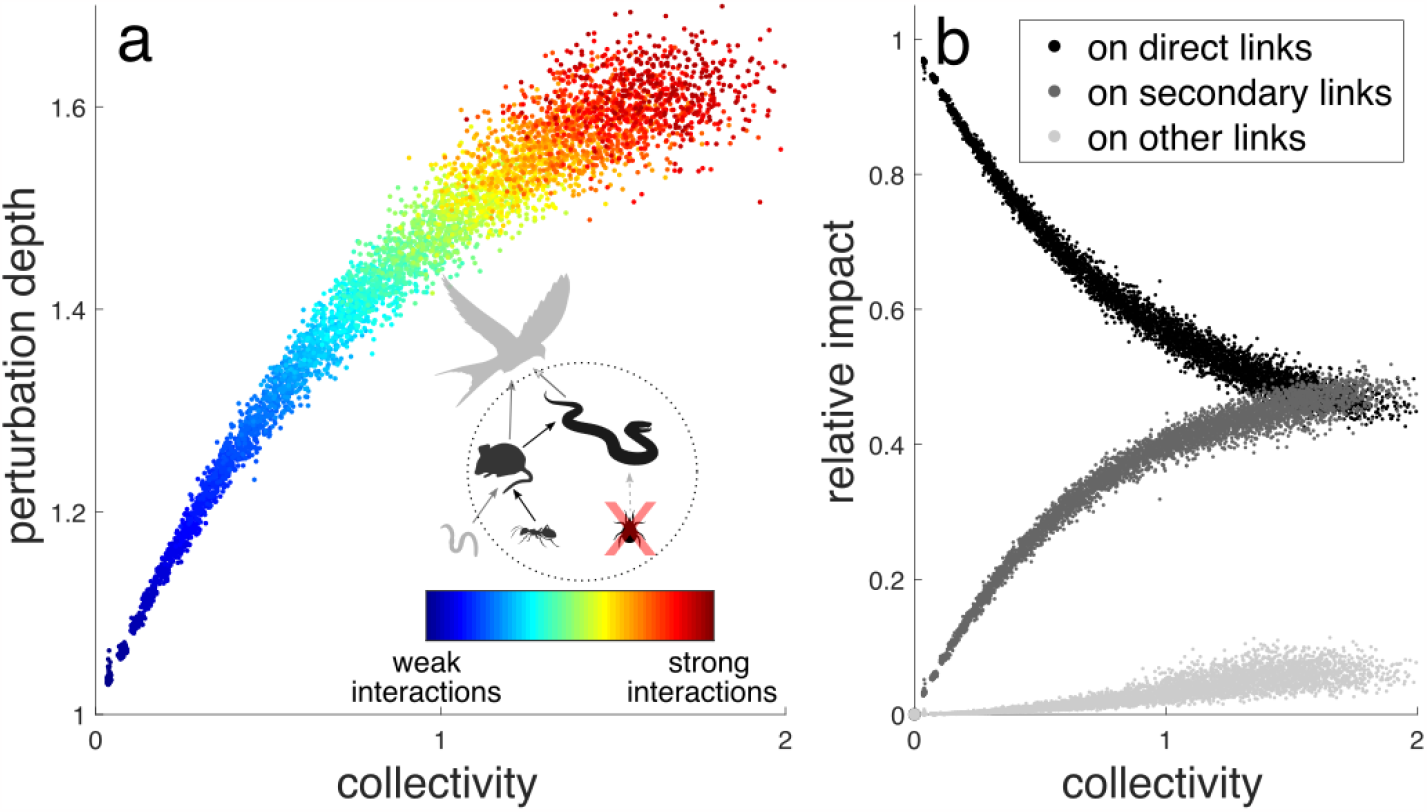
Perturbation depth and collectivity. For various communities, the effect of removing a single species is seen across the community. Left panel shows the perturbation depth, a measure of how deep into the network of species interactions does the perturbation reach. Right panel shows the average effect on the species in the community (all except the one species removed), partitioned into three groups: black for species directly interacting with removed species, dark gray for species directly interacting with the species in the black group, light gray for all other species. As collectivity increases the average effect on a given species becomes equal, regardless of its grouping (i.e. its position in the community structure), and therefore the perturbation depth increases (i.e. the effect of the perturbation if felt throughout the community). See Appendix S6 for more details.

### Temporal unpredictability

Indirect interactions between species require time to take effect. Thus, collective integration is expected to leave a signature in the relationship between short- and long-term response to a perturbation. Following a persistent change in abiotic conditions, a given species’ population will first react to the induced change in its intrinsic growth rate. Later, direct interactions between species will then take effect, followed by longer interaction pathways. However, if the strength of indirect interactions rapidly decays with their length, the latter will not substantially change the population dynamics; the long term outcome could have been extrapolated from the short term response. Therefore, the more collectively integrated the community, the less predictable the long-term response of a species should be.

We test this idea by randomly perturbing the intrinsic growth rates of all species of a community at equilibrium. We then measure the correlation between a vector of short-term response extrapolation *R*_*S*_ and a vector of the actual long-term response *R*_*L*_ (see Appendix S6), and define temporal unpredictability as the complement of that correlation:

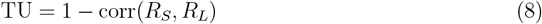

In Fig. 3 we compare this notion to collectivity *ϕ*. We see that the two are closely related, with unpredictability increasing steadily as collectivity grows, reflecting the fact that trajectories can, as indirect interactions come into play, change tendencies through time (right panel). Here too there is a caveat. If direct interactions are mediated by slow latent variables, such as unobserved species or modified environmental variables, time and length of interaction pathways need not be related. Collectivity thus leaves a unambiguous signature in temporal trends only if a separation of time scales exists between the factors that mediate direct interactions, and the actual population dynamics.

**Figure 3.**
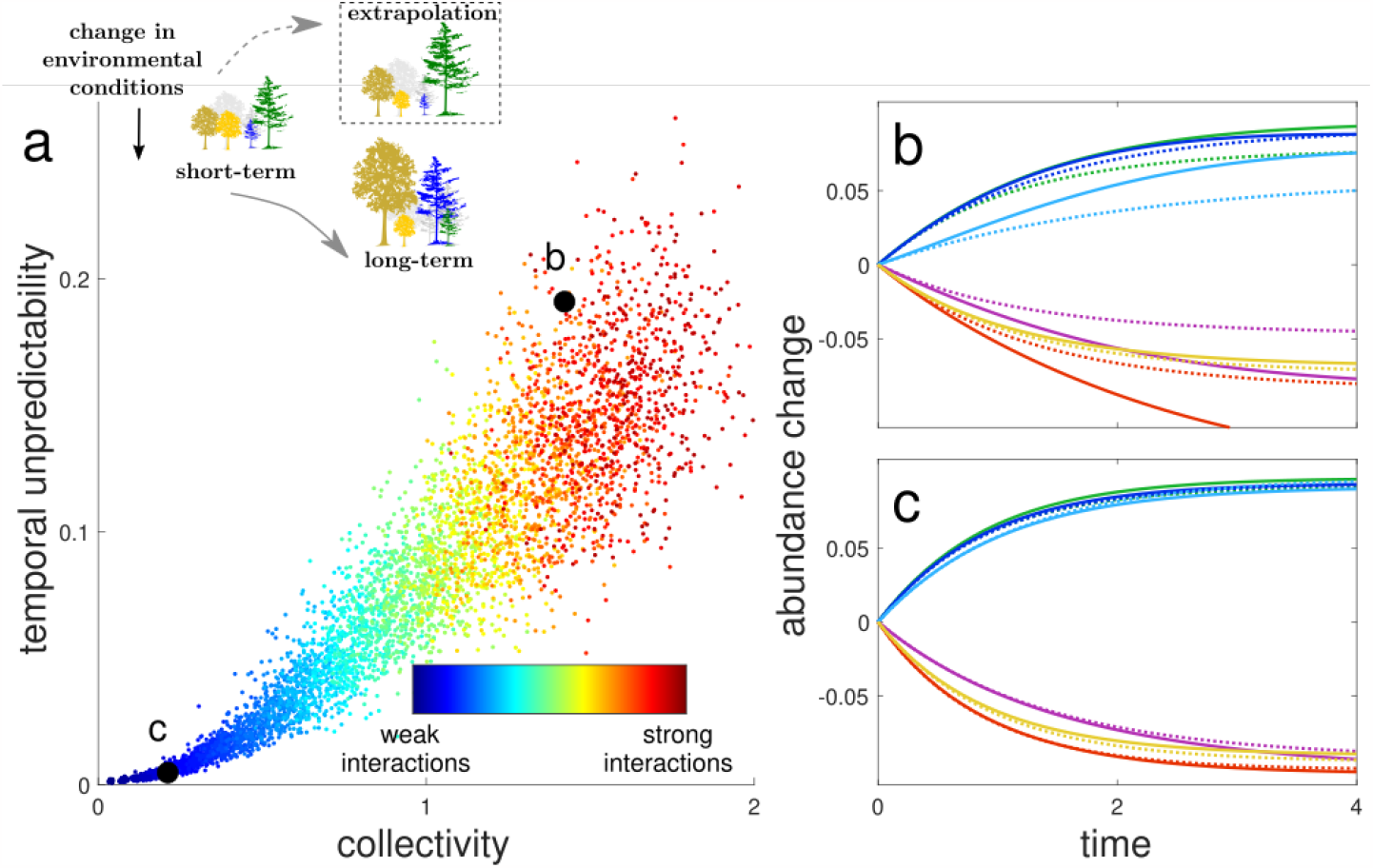
Temporal unpredictability between short-term and long-term response to perturbation. For various communities, the ability to predict the long-term response of a community to a perturbation from its short-term response is evaluated and shown. Left panel shows the temporal unpredictability, which gives a score of 0 for a perfect correspondence between short-term and long-term response. Black circles show the collectivity and temporal unpredictability for two communities, with the corresponding dynamics shown in the right panels. Right panels show the change in abundance for 6 species in each community, where the dashed lines show the extrapolated dynamics based on the short term fit (using the first 0 to 0.5 time units), whereas the solid lines show the actual dynamics. With higher collectivity the long-term behavior becomes less predictable, see Appendix S6 for more details.

### Biotic contribution to the realized niche

If species do not interact, only the abiotic environment (i.e. what cannot be attributed to the rest of the community) determines the species’ growth and abundance. In general, however, species change the environmental conditions perceived by other species. Intuitively, we can expect that the stronger the collective integration of the community, the more important and intricate this biotic contribution becomes (Levine *et al*., 2017).

To quantify this collective contribution, we start from the relative yield of a given species, *η*_*i*_ = *N*_*i*_*/K*_*i*_. This amounts to comparing mono-cultures to polycultures (second column of Fig. 4). The absolute difference between relative yield and unity is a measure of the net effect, on species *i*, of the biotic environment. We measure this effect integrated over the community and to make the result comparable across communities, we normalize by the sum of relative yields. This defines a measure of the Biotic Contribution to species realized niches:

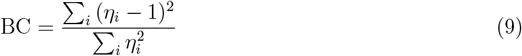

which is similar to the relative Euclidean distance ||*N−K*|| */*|| *N*|| between the realized community state *N* (expressing the realized niche) and what it would have been without interactions, *K* (the fundamental niche).

**Figure 4.**
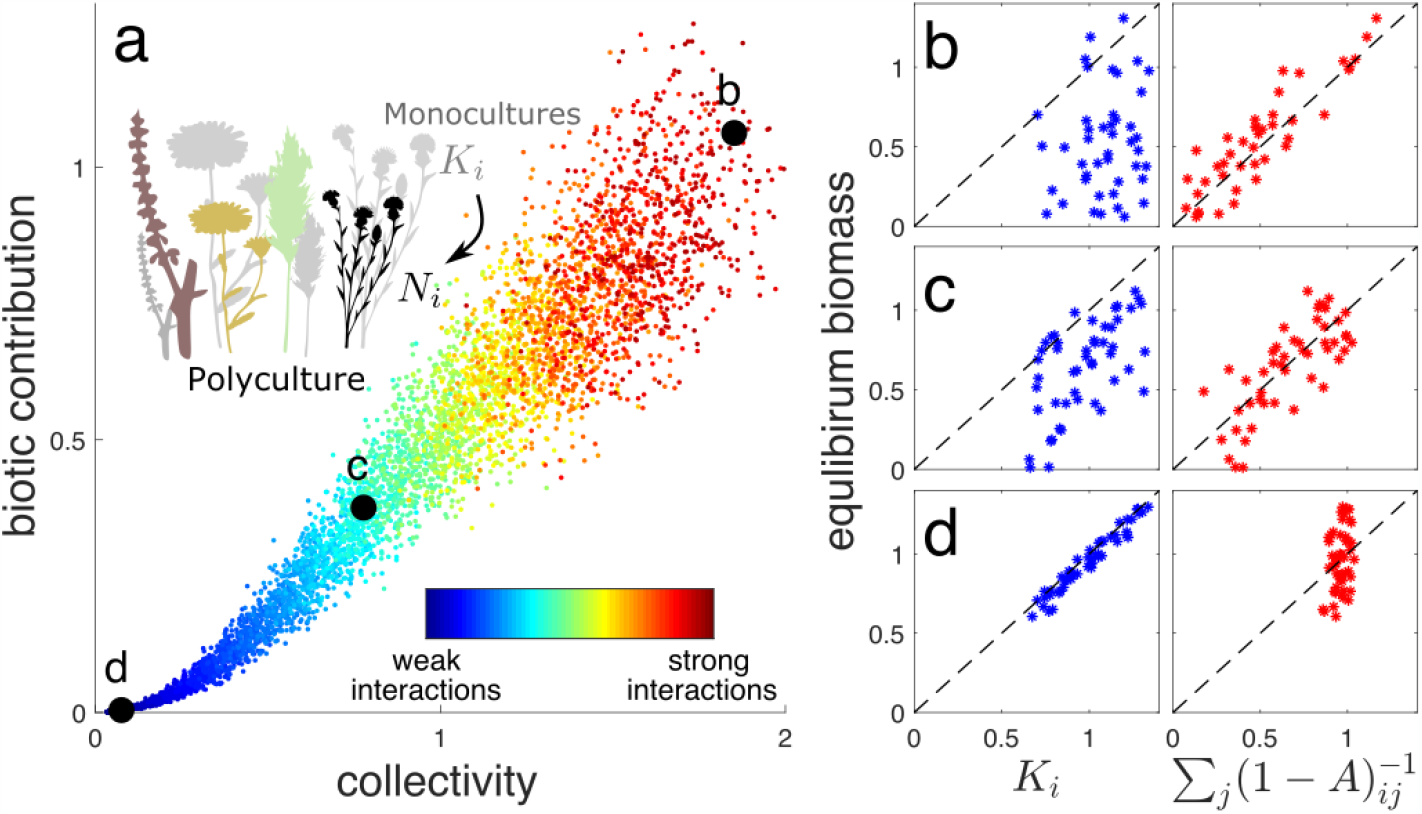
Biotic Contribution to species realized niches. The determinants of the community’s species abundance at equilibrium are evaluated. (a) the biotic contribution to the realized niche (eq. 9), with black circles highlighting several communities that are considered in the right panels. (b)-(d) Species equilibrium abundance for different communities (corresponding to black circles in panel a), compared with its carrying capacity (left in blue), or by contrast, with its abundance if all species had the same carrying capacities, so that differences in abundances are caused by species interactions only (on the right in red). Dashed black line shows the 1:1 line. See Appendix S6 for more details.

We can also characterise the raw contribution of the biotic environment by comparing the realized abundance of species to those achieved if species had the same carrying capacity (third column of Fig. 4). In fact, this amounts to asking how much a species abundance is explained by its *centrality* in the interaction network (Sharkey, 2017).

In the first panel of Fig. 4, we see that the collectivity parameter and the strength of the biotic niche Eq. (9), closely follow one another. For communities with collectivity parameter close to or larger than 1, species abundances are not at all explained by the abiotic environment, but instead are almost entirely controlled by the biotic environment set by the whole community. The caveat here is the requirement of a notion of carrying capacity, which makes senses for, say, plants but is ill-defined when considering consumer species.

### Collectivity, complexity and stability

In this section we relate collectivity to the *complexity* measure of May (1973), which we then use to assess the role of network structure, and discuss the sensitivity of collectivity to uncertain knowledge of biotic interactions. We will also clarify similarities and differences between collectivity and the widely studied notion of asymptotic *stability*.

We first invoke known properties of the spectral radius to deduce useful bounds on collectivity. Start from the fact that *ϕ*≤ || *A*||, where ||*A*|| is the spectral-norm of *A*, the maximal amplification of vectors’ length that this matrix can achieve (Reed *et al*., 1972; Trefethen & Embree, 2020). The spectral-norm is related to a simpler norm, via the following general equivalence relation (Reed *et al*., 1972):

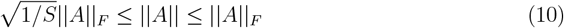

where ||*A*||_*F*_ is the FrobeniuIs norm of *A*: the square root of the sum of squared elements of *A*. Thus, 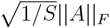 is really 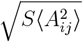, with 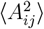 the mean of squared interactions (accounting for the trivial diagonal terms *A*_*ii*_ ≡0). We can develop this term a little more by introducing the network’s connectance 0≤*p*≤1 (i.e. the proportion of realized links) and *ξ*^2^ the second moment of interactions between connected species, so that

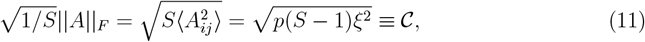

in which we recognize 𝒞, the complexity measure introduced by May (1972) in his seminal work on Random Matrix Theory and the complexity-stability debate (McCann, 2000).

Finally, given that *A*_*ii*_ ≡ 0 (see Appendix S1), we can transform the general equivalence relationship Eq. (10) to deduce a similar relationship between complexity and the spectral radius of interaction matrices:

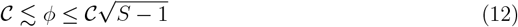

The upper bound is sharp^2^ while the lower bound is an approximate one (i.e. sharp and impossible to cross only for normal matrices (Trefethen & Embree, 2020; Asllani *et al*., 2018)). The bounds on *ϕ*, because they do not depend on the way interactions are actually distributed, can be used to quantify the role of structure. Rescaling *ϕ* relative to the upper and lower bounds, we can define a measure for which a value of 1 means that the upper bound is reached, indicating that structure maximizes the spectral radius, thus collectivity. A value of zero would imply that *ϕ* is unaffected by randomizing interaction terms. Because the lower bound is sharp only for normal matrices, values can become negative if the structure allows it. In Appendix S2 we provide examples of structures, from competitive networks to food-web models, that greatly affect collectivity, allowing to move from one bound to the other.

The approximate lower bound should thus be seen as the collectivity that ‘comes for free’ if the matrix was random, recalling that non-random structure could reduce collectivity far bellow this baseline. This baseline expectation can be refined to account for additional statistical features of interactions, such as their mean, variance and reciprocity (the correlation between *A*_*ij*_ and *A*_*ji*_, which is negative in food webs, and typically positive in competitive networks). The complete description of the spectra of large random matrices given by Allesina & Tang (2012), then gives a random matrix prediction for *ϕ* (see Appendix S2). The accuracy of this prediction, which we test in Fig. S3, demonstrates that, in the absence of a clear structure, collectivity can be reliably estimated using only summary statistics of interactions. In Appendix S2 we test this claim by estimating collectivity from incomplete knowledge of interactions. Concretely, we assume that only a fraction of links are known, while the rest is inferred based on the statistics of the measured interactions. In the absence of structure this procedure works well. Any method that can infer interaction statistics, without necessarily providing any reliable information about individual interaction terms, would still give an accurate approximation of collectivity. However, if an underling structure exists in the network, so that collectivity is far from the baseline expectation, basic statistics of interaction strength are insufficient. They must be combined with knowledge of structural features (Asllani *et al*., 2018). In Appendix S2 we illustrate this idea using a cascade food-web model (Pimm *et al*., 1991), where predators feed on smaller species. The trophic structure generates a substantial departure from the random baseline. However, given basic features of this structure, only summary statistics of attack rate and trophic efficiency are required to accurately predict the food-web’s degree of collectivity.

If *ϕ* is associated with May’s complexity measure, should it not be directly related to stability, at least sensu May (1972)? The linear stability criterion is that all eigenvalues of the Jacobian matrix at the steady-state must have a negative real part (Lyapunov, 1892). This is essentially^3^ equivalent to all eigenvalues of *A* having a *real part* smaller than 1 (Gibbs *et al*., 2018). Collectivity *ϕ* is instead the largest eigenvalue *modulus*. As is made clear once represented graphically (see middle inset of Fig. 5), if the system is unstable, *ϕ* is necessarily larger than 1. However, even if *ϕ* is large, the real part of the associated eigenvalue can still be arbitrarily small (this is the case in the consumer-resource model of eq. 6). So instability implies a high degree of collective integration *but the converse is not true*. Large collective integration does not imply instability (right panel of Fig. 5).

**Figure 5.**
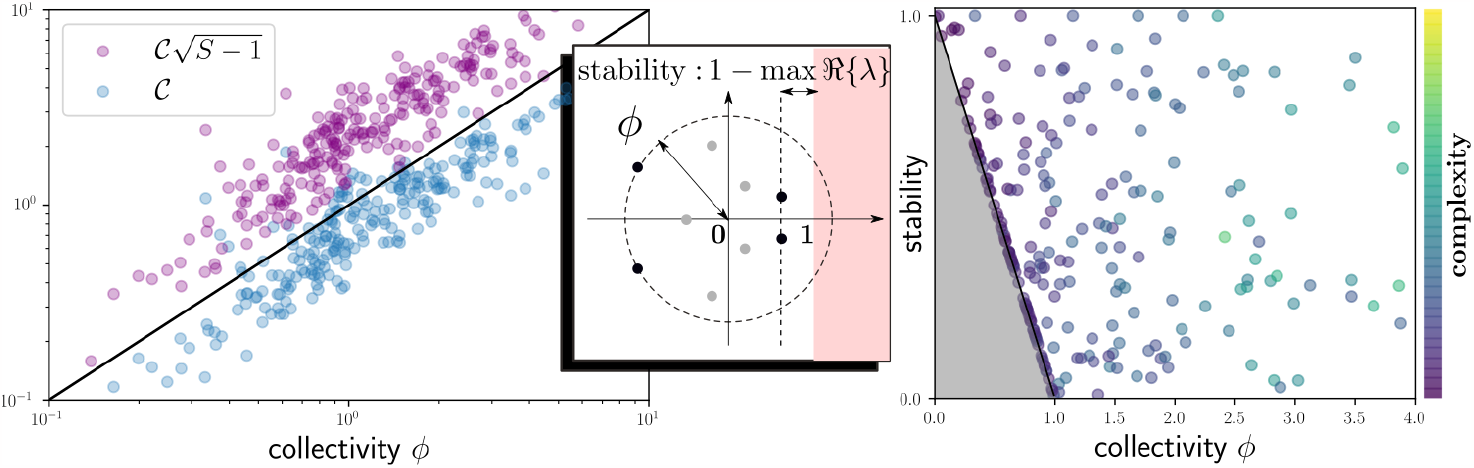
Left: May’s complexity 𝒞 sets the bounds of collectivity, illustrated here for 242 random stable matrices (out of 1000 generated) of size 5 *×* 5 to 15 *×* 15, representing interaction matrices *A* of various connectance, interaction strength, variance, and pair-wise symmetry (notice the log-log scales). In purple the sharp upper bound 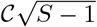 where *S* is the size of the community (purple points are always above the one:one line in black). In blue the approximate lower bound𝒞 (blue points mostly remain below the one:one line). **Middle inset:** Stability is here a distance to instability 1*−*max ℜ{*λ*}, where max ℜ{*λ*} is the maximal real part over all eigenvalues *λ*∈ℂ of the interaction matrix. If this real part attains 1, the community is unlikely to be stable –shaded region is the instability domain. Collectivity is instead the radius of the smallest disc centered on 0 that contains all eigenvalues. **Right: collectivity and (in)stability are not equivalent**. Same matrices as on the left panel. We see that large values of collectivity are allowed even if we restrict to stable systems. The edge of the grey region represents *y* = 1 − *x* which is what would be expected if collectivity and stability where associated to the same eigenvalue.

## Discussion

In this theoretical piece, we have addressed two interlinked issues, which are both of conceptual and practical importance: i) how to bridge between reductionist and holistic conceptions of ecological communities, with a quantitative well defined axis, and ii) how to better understand and quantify the dynamical role of interaction structure. These two points go back to two classic dichotomies of ecology, reductionist vs holistic views and dynamics vs structure. They are also practical. Point i) clarifies when we can hope to predict behavior using incremental causality. Point ii) helps us understand community dynamics even with limited data.

The premise of our work, is that, in nature, dynamical inter-dependency between species occurs not only though direct interactions, via predation, facilitation or competition, but through potentially much longer indirect interaction pathways (Wootton, 1994). When long indirect pathways significantly contribute, over relevant time-scales, to population dynamics, the state of a species can depend on many -if not all-other species in the community. We hypothesise that in such instances, the incremental causal thinking that characterises individualistic approaches would become misleading (Levins, 1974). We also expect that when contributing interaction pathways are long, perturbations propagate further; short-term and long term responses of species become uncorrelated; and finally, that in such collectively integrated communities, it is the biotic environment (and not the abiotic one) that shapes abundance distribution patterns.

To make progress on these ideas, we studied the spectral radius *ϕ* of a community’s interaction matrix, i.e. its largest eigenvalue modulus. We showed why it is this precise feature that determines the length of non-negligeable indirect interactions in dynamical community models. Moreover, we explained that when *ϕ >* 1, arbitrary long interaction pathways can have as much importance as shorter ones. In such instances, it becomes impossible to decompose long-term effects as a sum of dominant direct or indirect interaction pathways. In this sense, incremental causality breaks down.

We proposed to see *ϕ* as a measure of collective integration, as it drives empirically relevant collective behaviour. On simulated ecological communities, we demonstrated that *ϕ* drives *Perturbation depth*, defined as the network distance covered by perturbations initially affecting only a single species. We also showed that *ϕ* increases *Temporal unpredictability*, defined as the discrepancy between species long-term behaviour after a change in environmental conditions, and the extrapolation of their dynamics based on their short-term response to the environmental change. We finally related the value of the collectivity parameter with a measure of *Biotic Contribution* to species realized niches, defined to quantify the degree to which abundances of species grown in monocultures, fails to predict the abundances of those same species when grown together.

We then showed that our measure of collectivity parameter is controlled by May (1972)’s complexity measure, a summary statistics of absolute interaction strength. Complexity, together with species richness, determines both an upper bound on collectivity and a baseline expectation. Thus complexity sets the baseline for collectivity, while the interaction structure determines how much it will depart from the baseline, either reducing collectivity or increasing it towards the sharp upper bound. We identified structures that push towards maximal collectivity: evenly distributing interactions do just that, as well as increasing pair-wise symmetry. By contrast, triangular (i.e. hierarchical) structures, because they buffer feedbacks (Tilman, 1994), drastically reduce collectivity. In Appendix S2 we consider a cascade food web model (Pimm *et al*., 1991), showing that trophic efficiency, which determines how predators utilize prey biomass, is a major structural driver, making food-webs go from low collectivity when efficiency is low, to near-maximal values for high efficiency

In a similar vein, (Staniczenko *et al*., 2013) used the spectral radius of adjacency matrices to quantify the nestedness of bi-partite interaction networks (plant-pollinator networks are often nested: specialist insects typically pollinate a subset of the plants pollinated by generalist species). In their work, only presence or absence of links are known, so the spectral radius should not be seen as a measure of collectivity, as the later depends on the quantitative value of interactions. Nonetheless, by comparing with the random baseline and the upper bound (Chung, 1997), we see which structures favour collective behaviour, those that have no effects, and those that ought to reduce it. Given that (Staniczenko *et al*., 2013) find that nestedness is associated with a higher than expected spectral radius, we may say that, *ceteris paribus*, nestedness of bi-partite networks favours collective behaviour.

Overall, this is a novel perspective on ecological network structure: instead of asking whether biotic structures tend to make communities more or less stable (May, 1972), we ask whether those structures tend to make communities more or less collectively integrated than if they had been randomly assembled.

In collectively integrated communities, long indirect pathways between species, whose importance unfolds over variable time scales, do not favour predictive power. As previously shown (Yodzis, 1988, 2000; Barabás *et al*., 2014), this is certainly true if one tries to predict the effect that a perturbation will have, in the long-term, on a given species when the latter is embedded in a complex ecosystem. But this typical sensitivity of individual variables need not imply that all properties of a community or ecosystem are sensitive and/or unpredictable (Daugaard *et al*., 2022). We know, in particular, that many aggregate features of complex models are robust to uncertainty in parameters (Barbier *et al*., 2018). From our work, we can predict when those long indirect interaction pathways cannot be ignored or even simply added up. In such instances we must change perspective, and focus our efforts towards robust ecosystem or community-level properties of the natural system under study (Goldford *et al*., 2018; Bergelson *et al*., 2021; Sanchez *et al*., 2023).

### Perspectives

Our measure of collective integration is defined on isolated community models, which are drastic simplification of real communities, that are open to migration, structured spatially, and neither deterministic nor stable (Hastings, 2004).

It is not obvious how to translate our mathematical analysis to transient, far-from equilibrium dynamics or draw conclusions on the role played by spatial structure and scale in determining the degree of collective integration of ecological systems. For the latter, this relates to the meta-community concept (Holyoak *et al*., 2005) and the quest to better understand the spatial scaling of ecological interactions (Gravel *et al*., 2016; Galiana *et al*., 2018, 2022). This is a promising direction, that could help better formulate the scale transitions of ecological patterns, as it is commonly thought that at large scales, ecological systems are mostly determined by abiotic drivers thus suggesting a negative collectivity-scale relationship. Considering transient states is also challenging. Powerful dynamical techniques do exist to tackle transients in complex systems (Roy *et al*., 2019). But even without abandoning the notions of stationarity or equilibria, we could expand our formalism to address the time-scale dependency of collectivity. A way forward may be the study of power-spectra of ecological time-series: decomposing the temporal fluctuations of populations over various times scales to infer the interaction structures that generate such signals (Krumbeck *et al*., 2021). Here we suspect that variations over longer time scale reflect the collective nature of communities more than those at much shorter time scales, as the latter would not allow for indirect interactions to manifest, as seen in Fig. 3.

Finally, can we measure the degree of collective integration in empirical data? The fact that collectivity is controlled by aggregate interaction statistics (see Fig. 5) means that we need not precisely measure all species interactions to estimate collectivity bounds. In fact, basic interaction statistics entirely determine collectivity in the absence of structure (Allesina & Tang, 2012), which can allow its estimation from incomplete or noisy interaction strength data (see S2). But more generally, we can expect that there will often exist a combination of basic aggregate features of structure and interaction strength that drive our measure of collectivity. For instance, in the food-web model example shown in S2, basic topological structure, mean attack rate and trophic efficiency are its main drivers. This is reassuring as any theoretical notion that is sensitive to fine scale details is guaranteed to be irrelevant. This being said, can we reliably estimate collectivity from, say, time-series analysis (Sugihara *et al*., 2012) or residual correlations in species distributions (Ovaskainen & Abrego, 2020). In Appendix S2, we give a proof of concept for the latter, showing that in unstructured competitive communities, collectivity is surprisingly well estimated from the correlation between species abundances across environments. Developing such ideas could be a realistic way forward towards quantifying the dynamical importance of biotic interactions in complex communities.

## Acknowledgements

ML and JFA were supported by the “Laboratoires d’Excellences (LABEX)” TULIP (ANR-10-LABX-41). NG received funding from the European Union’s Horizon 2020 research and innovation programme under the Marie Sk∤odowska-Curie grant agreement BIOFOODWEB (No 101025471). The authors thank two anonymous referees for their thought-provoking critiques that greatly help improve the manuscript. JFA thanks Mathew Leibold, Sonia Kefi and members of the Biodicee team, Leo Ledru as well as the whole INTP community for their encouraging and thoughtful comments during the writing of this manuscript.

## Box 1

**direct, net and indirect interactions in stable community models**

Consider a model that specifies the growth rate *g*_*i*_ of all species, as a function of their joint abundances ***N*** = (*N*_*i*_) :

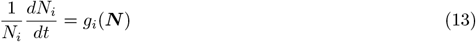

In Generalized Lotka-Volterra (GLV) models *g*_*i*_(***N***) = *r*_*i*_ +∑_*j*_ *a*_*ij*_ *N*_*j*_, with *a* = (*a*_*ij*_) representing percapita interactions and ***r*** = (*r*_*i*_) the vector of species intrinsic growth rates. We assume that the community is in a steady state ***N*** ^*\**^, so that *g*_*i*_(***N*** ^*\**^) = 0. In that state, we want to define direct and net species interactions, relate them to one-another, and show *in what sense net interactions emerge as a sum of indirect* ones, a claim that we use to quantify the collective integration of the community.

### Direct interactions

reflect the sensitivity of the growth rate of one species, to a change in abundance of another. In mathematical terms, this amounts to defining the matrix of partial derivatives

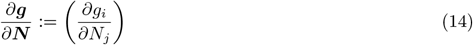

which for GLV models, coincides with the matrix *a* = (*a*_*ij*_).

### Net interactions

*are the reciprocal of direct interactions*: the long-term sensitivity of the abundance of a species, to a permanent shift in the growth rate of another (Novak *et al*., 2016; Montoya *et al*., 2009). In matrix form, net interactions can be written as

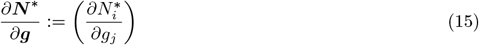

To clarify the meaning of Eq. (15), imagine applying a small press perturbation *δ****g*** on the species’ growth rates. In contrast with the way direct interactions are defined, we now *let community dynamics play out*, ultimately leading to a shift in equilibrium abundances *δ****N*** ^*\**^, so that

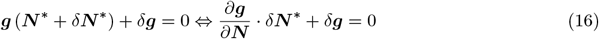

This expression can be inverted to show that the matrix of net interactions is indeed the inverse of the matrix of direct interactions:

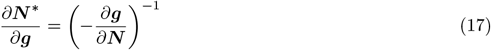

In GLV models this matrix also determines the steady state ***N*** ^*\**^ = (*−a*^*−*1^) *·* ***r***.

### Indirect interactions and the collectivity parameter

Direct and net interactions have reciprocal units. Furthermore, if we multiply the direct interaction between species *i* and *j*, with the direct interaction between species *j* and *k* this would change dimensions, and define an indirect interaction between species *i* and *k* that cannot be compared to neither direct nor net interactions. However, by defining direct interactions *relatively to self-regulation*, defined for any species *i* as 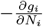, we can revisit the connection between direct and net interactions such that a coherent notion of indirect interaction emerges. Let direct, *non-dimensional* interactions be

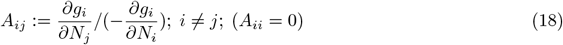

In GLV models this corresponds to *A*_*ij*_ := *a*_*ij*_*/*(*−a*_*ii*_). If *D* is the diagonal matrix encoding species self-regulation, and 𝕀 the identity matrix, direct interactions can be written as

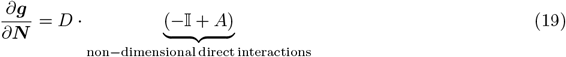

which indeed have the same units as *D*. From Eq.(17), it follows that

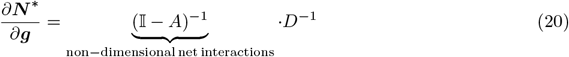

If the elements of *A* are small, the *non-dimensional* net interaction matrix (𝕀*− A*)^*−*1^ can then be written as a convergent infinite series

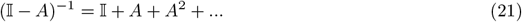

This series enables us to define **indirect interactions** of order *k* as the elements of *A*^*k*^. Indeed (*A*^*k*^)_*ij*_ is the sum of all non-dimensional interaction pathways of length *k* that lead from species *j* to *i* (allowing for loops), drawn in the interaction network. Those should not be confused with higher-order interactions (Battiston *et al*., 2021), which are defined as non-linear interaction terms in community models (See discussion in Appendix S4). It is in the precise sense of Eq.(21) that net interactions emerge as a sum of indirect ones. Our measure of collectivity *ϕ* is the spectral radius of *A* (Trefethen & Embree, 2020), namely:

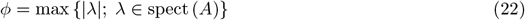

which controls the rate of convergence of the series Eq.(21), and thus **that contribute to net interactions**.

## Supporting Information

In this Supplementary Information, we give further detail about collectivity and its estimation, as well as to other quantities that relate to it, including the signatures described in the main text. In S1 we give further details about the relation of collectivity with two well known notions, stability and complexity. In S2 we discuss ways of estimating collectivity and their limitations. In S3 we shortly discuss the definition of a community, while in S4 we discuss the differences between indirect and higher-order interactions. In S5 we detail how the communities that were used for the main figures were assembled, and in S6 we give further detail about the signatures of collectivity.

## S1 Collectivity, complexity and stability

To quantify the importance of indirect interaction pathways in community dynamics, we reduced the problem (see Box 1 in the main text) to a study of the inverse of the matrix of relative interactions *−*𝕀 + *A*, which has the remarkable property to provide an analytical extension to an infinite series of indirect interaction pathways:

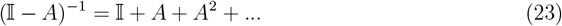

The left term, the matrix inversion, formalizes long-term ecological dynamics caused by interactions between species. The convergence of this series is determined by the spectral radius *ϕ* of *A*, the maximal eigenvalue modulus. The general bounds on *ϕ* allow us to connect our work with May’s complexity measure 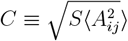, via the (approximate) chain of inequalities

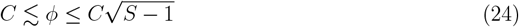

The upper bound is sharp, but the lower bound only holds for normal matrices (triangular or quasi-triangular interaction structures can lead to a much smaller spectral radius). This chain can be used to assess the role of interaction structure, given by the position of *ϕ* within those bounds, a way to control for interaction strength and connectivity.

Importantly, the term (*I−A*)^*−*1^ in Eq. (23) can still be meaningful even if the series diverges, so even when *ϕ*≥1. The divergence of the indirect interaction series only reflects an instability in the underlying model when the spectral radius is also the maximal real part of the eigenvalues of *A*. Thus there can be divergence of the series while the dynamical system associated to *A* remains stable.

### S1.1 Demonstration of how high collectivity does not lead to instability

As noted in the main text, although collectivity may seem related to the notion of linear stability (as they both depend on spectral properties of a matrix representing the community), they are quite distinct. In fact, at high collectivity the series of indirect interactions can diverge without implying unstable behaviour, but may still lead to counter-intuitive phenomena.

If not instability, surely something else must be happening to the dynamics when *ϕ >* 1. What we now show is that the divergence of the indirect interaction series allows for counter-intuitive phenomena, unforeseen when only considering short indirect interaction pathways. To do so we give a simple yet important food-web example, that can maximize collectivity over anti-symmetric interaction networks 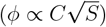. In this example, the counter-intuitive phenomenon that occurs when *ϕ >* 1 is that long indirect interaction pathways make net interactions between prey and predator *decline* as attack rate *increases*, so that direct and net interactions become negatively correlated.

The model is the following: a single predator feeds on *S−*1 preys, with an attack rate *a >* 0 which is converted into growth rate with efficiency 0 *<* ϵ ≤1 (see Fig. S1). The matrix associated to this food-web reads

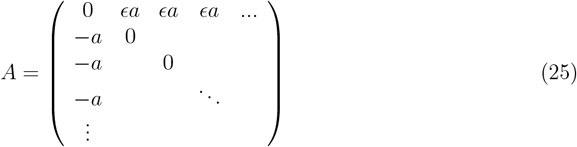

**Figure S1:**
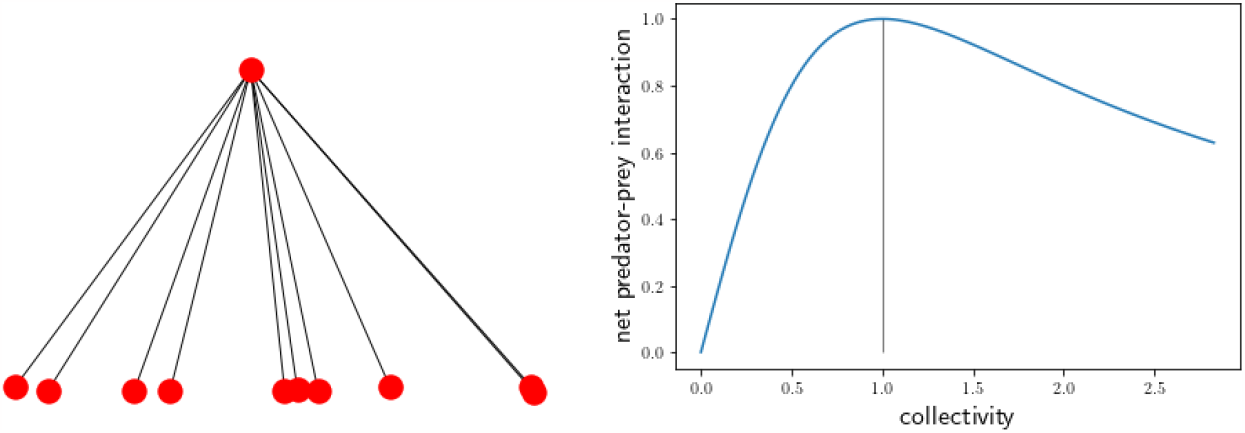
At low collectivity, net interactions grow with collectivity *ϕ*, which is proportional to attack rate, but beyond *ϕ* = 1, a non-intuitive phenomenon occurs that is caused by arbitrarily long indirect interaction pathways: net interactions start to decay, despite larger and larger attack rates. In such systems, the dominant real part of the spectrum is unaffected by parameters, and remains 0.

The complexity *C* of this interaction network is

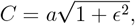

and its eigenvalues are purely imaginary, with a single non zero pair that therefore gives the spectral radius *ϕ*:

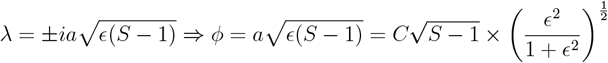

thus approaching the general upper bound 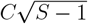on collectivity as *ϵ∼*1, while being potentially arbitrary small if *ϵ∼*0. In this example stability is unaffected by the parameters as the imaginary part of eigenvalues remains 0. Yet collectivity *ϕ* can be much larger than one (even if attack rate is small). Let us now compute indirect interactions between the predator and a prey, to see how their sum behaves. Recall that, when the sum converges it coincides with the net interaction between the predator and a prey. First, by definition, the direct interaction between the predator and one of its preys is the attack rate *−a*. Obviously there is single direct path:

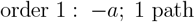

The second order term is 0, as no pathway of this length connects the two species. Third order interactions correspond to a loop that goes through any prey before going back to the focal one. There are *S −* 1 such paths:

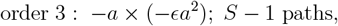

We can go on and show that indirect interactions of order 2*n* + 1 read

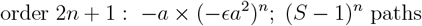

Net interactions are the sum all these indirect ones, and can therefore be expressed as the series

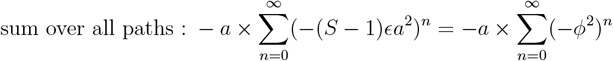

Where we could recognize the spectral radius *ϕ*. We see that when the attack rate *a* reaches the critical value

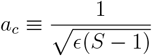

Then *ϕ* = 1 and the indirect interaction series is an infinite alternate sum of 1 and*−*1, which consequently does not converge, as expected from our theory. So what happens then?

As stated above, the eigenvalues of *A* are strictly imaginary: the system’s stability is unaffected. Instead, the divergence of the above series signals a counter intuitive behaviour of net interactions between predator and prey.

To see this we first make an analytical continuation of the series. This is easy to do because, for *ϕ <* 1, we have that

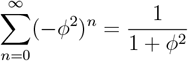

where the right-hand side extends the series beyond its domain of convergence. Thus, for all values of *ϕ*, the net interactions between prey and predator are actually finite and read

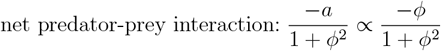

We see here that net interactions between prey and predator will start to *decline* when *ϕ >* 1 (see Fig. S1 above). Due to arbitrarily long indirect interaction pathways net interactions become anti-correlated with direct interactions (i.e attack rate), a phenomenon that cannot be understood when considering only a few, albeit long, indirect interaction pathways.

### S1.2 Spectral radius and Frobenius norm of zero-trace matrices

Here we provide more details on the spectrum of zero-trace matrices, with the goal of corroborating the upper-bound on collectivity 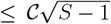, stated in the main text. Note that this bound is only a slight improvement on 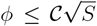 which is immediately deduced from *ϕ ≤* ||*A*|| *≤* ||*A*||_*F*_, and 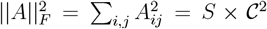 (by definition of 𝒞). We that,because *A* has zero-diagonal (thus zero-trace), we can shrink the bound *ϕ ≤* ||*A*||_*F*_ into 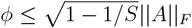. This enables us to claim that uniform competition amongst all species in a community, for which it is easy to show that 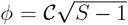, is the interaction structure that maximizes collectivity. If we didn’t known that this was the true upper bound on *ϕ*, we would have to speculate on the existence of other interaction structures that could increase collectivity further.

**Figure S2:**
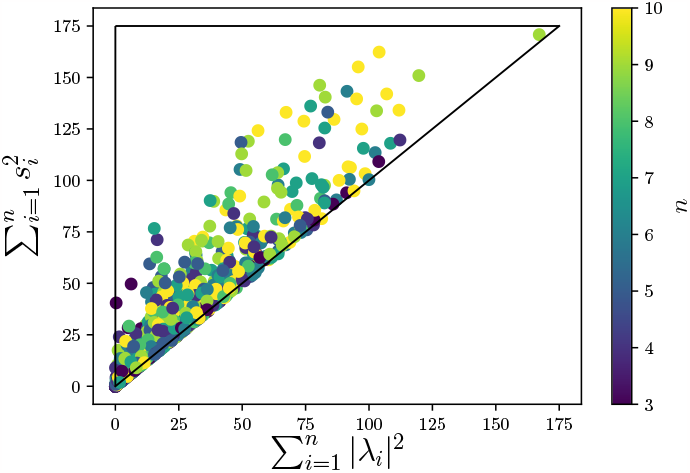
Test of conjecture 4: for a wide array of randomly generated matrices of size *n × n* with *n* = 3, …, 10. We could never find any matrix for which the inequality 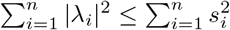did not hold (and we know that it always holds for *n* = 2).

#### Lemma 1.

*Let A ∈ M*_*n*_ (ℝ) *(a real square matrix) such that TrA* = 0. *Let λ*_0_, *λ*_1_, …*λ*_*n−*1_ ∈ *ℂ, i* = 0, …, *n−* 1 *its n eigenvalues (possibly repeated), ordered in decreasing order of modulus, so* |*λ*_0_| *≥* |*λ*_1_| *≥* … *≥* |*λ*_*n−*1_|. *It then holds that*

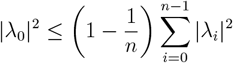

*Proof*. If *λ*_0_ has an imaginary part, then it comes as a conjugate pair with 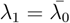, and the two have equal modulus. In this case 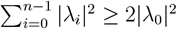. Note that that 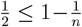 for all *n ≥* 2, so in this case the upper bound holds. Now suppose *λ*_0_ is real. Due to the fact that Tr*A* = 0, and that *A* is real, we have 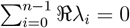. The maximal configuration for the rest of eigenvalues is that they all have the same real part, equal to 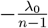. In this case 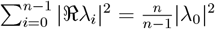.This shows that, when *λ*_0_ is real, it always hold that 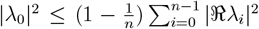 which is obviously smaller that 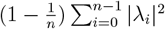.

#### Definition 2.

Let 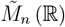 be the non-empty subset of *M*_*n*_ (ℝ) comprised of matrices *A* such that 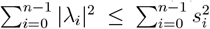, where 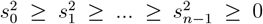 are the eigenvalues of the positive-definite matrix *A*^*T*^*A*.

For normal matrices, |*λ*_*i*_| = *s*_*i*_, 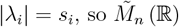 contains normal matrices.

#### Lemma 3.

*For any* 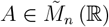 *such that TrA* = 0, *it holds that*

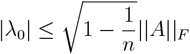

*where* 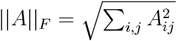 *is the Frobenius norm of A*.

*Proof*. From lemma 1 we have that 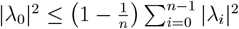 and if the matrix is in 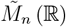 then it follows that 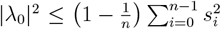. On the other hand, we may note that 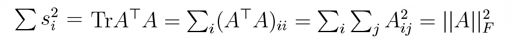. □

**Conjecture 4**. 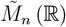*is a generic set. This means that, if there exists a real square matrix that is not in* 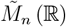, *then arbitrarily small perturbations of this matrix will be* 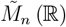.

We can show by a direct computation that 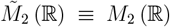. In higher dimensions extensive simulations (some shown in figure S2) support the idea that it is very unlikely that a given real matrix does not belong to 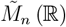.

## S2 Estimation of Collectivity

### S2.1 Use of Random Matrix Theory to estimate the collectivity parameter

We can apply the work of Allesina & Tang (2012) on the stability of large random ecosystem models to give a random matrix theory (RMT) prediction of the spectral radius of large matrices. Because RMT predictions only require summary statistics of matrices, we can use this result to explain why, in the absence of clear interaction structure, the collectivity parameter can be estimated without precise knowledge of all interaction strength between species of a community.

If *μ* and *σ*^2^ are the interaction mean and variance, *p* the connectance and *γ* the reciprocity of interactions (e.g. negative for predator-prey, positive for competition), then RMT predicts that

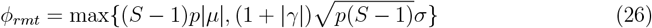

This prediction works well, even for sparse matrices of reasonable dimensions (see Fig. S3). In fact, we can use this result to answer how empirically accessible our collectivity notion is. For unstructured systems, an estimation of basic network statistics such as connectance, mean, variance and reciprocity of interactions will allow a robust estimation of collectivity. This explains why, when considering large enough randomly generated interaction matrices, extrapolating unknown interactions based of a faction of observed ones is enough to reliably estimate the spectral radius (Fig. S4).

### S2.2 The impact of interaction structure on the estimation of collectivity from incomplete data

When an underling structure exists in the network, basic statistics of interaction strength can fail to predict the spectral radius. This means that, when dealing with noisy or incomplete data, one cannot naively extrapolate unknown interaction coefficients to infer the spectral radius. This does not imply that it is inaccessible, but rather that structural knowledge about the network is necessary (Asllani *et al*., 2018).

To illustrate this point, consider a cascade food-web model (Pimm *et al*., 1991), where species are organized along a niche axis (such as body size) and can feed, with some probability *p*, on any species below them. For each predator-prey pair, we draw the attack rate between predator and prey as a positive random variable *X* (taking values say, between 0 and 2*a*) and define the interaction pair as *−X* and *ϵX*, where *ϵ ≤* 1 represents trophic efficiency. As the latter varies from 0 to 1 the interaction matrix goes from being triangular to perfectly anti-symmetrical. To see how this structure affects collectivity we rescale the latter with respect to the general bounds. A value of 100 means that the upper bound is reached. A value of zero means that the lower bound is reached. Because the lower bound is only an approximation, sharp only for normal matrices, values can become negative if the structure allows it (in the example collectivity can be made arbitrarily small by reducing trophic efficiency). Suppose that along a gradient of trophic efficiency, we want to estimate the collectivity of such a food-web. We see in Fig. S5 that the RMT prediction (in black) that neglects structure does not capture, even qualitatively, the major structural effect that trophic efficiency has on the collectivity of the food-web. Yet values of collectivity of food-webs that only share the same basic structure and attack rate statistics remain similar. Because estimations differ in terms of the random realization of trophic links and strength, we deduce that if we know the basic structure of the network and can estimate trophic efficiency, the collectivity parameter can then be reliably estimated without precise knowledge about interaction links and strengths.

**Figure S3:**
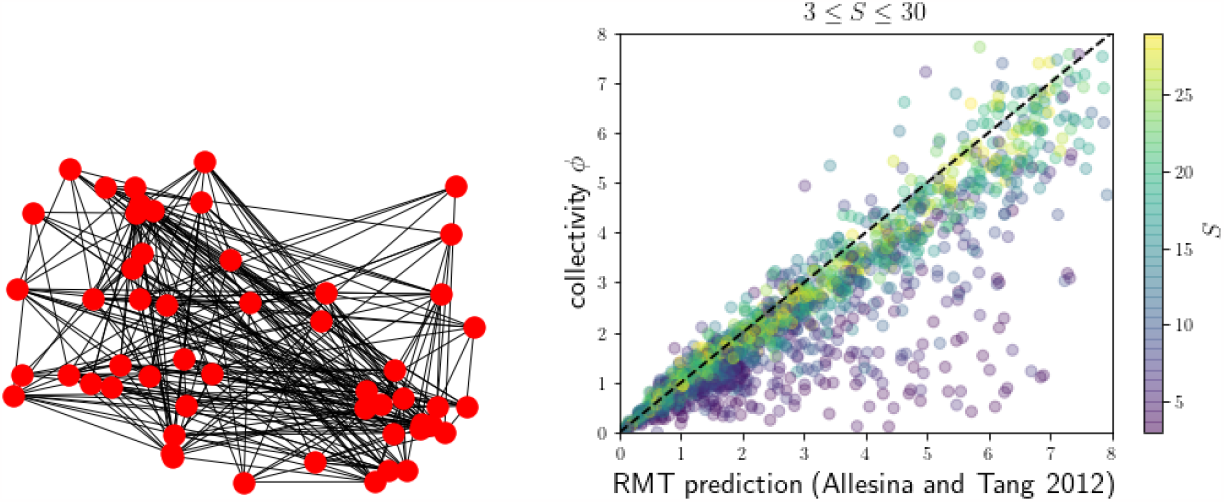
RMT prediction for the spectral radius of stable random matrices, of size 3*×*3 to 30*×*30 with randomly drawn connectance, interaction strength moments and reciprocity, colors correspond to the size of the matrix. Because RMT relies on large matrices, warm colored points, corresponding to larger matrices, cluster on the one:one line.

### S2.3 Inference from correlation data (proof of concept)

Here we provide a proof of concept for the robust estimation of collectivity using species abundance correlation data. Suppose a community is observed at different times, or different locations, so that stochastic noise or environmental variations, generate a variance in species abundances. Can we, at least in principle, extract information about species interactions based on the correlation between species? Here we show that even we cannot reliably estimate specific pair-wise interactions, there is still hope to deduce collectivity. Consider a GLV model, with a fixed interaction matrix *A*, where we model environmental differences between many patches as small variations *δK* in species’ carrying capacities (caused, e.g., by variations in growth rates). At equilibrium, the ensuing abundance variations will then read *δN* = (*I − A*)^*−*1^*δK*. The abundance covariance matrix across patches that ensues reads

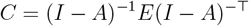

where *E* is the environmental covariance matrix across patches. The correlation matrix *ρ* is *R* = *D*^*−*1*/*2^*CD*^*−*1*/*2^, where *D* = diag(*C*). We see that *ρ*, which is in principle observable, explicitly depends on interactions *A*, that we would like to infer. Yet because *ρ* is symmetrical and bounded, and also depends on the environmental matrix *E*, we can guess that we cannot always simply invert the function *ρ*(*A, E*) = *ρ* to get *A*. There are some particular cases where we can however, which we can use as an ansatz for the general case^4^. In those favourable situations, after a few lines of matrix algebra we get to

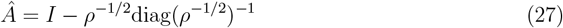

**Figure S4:**
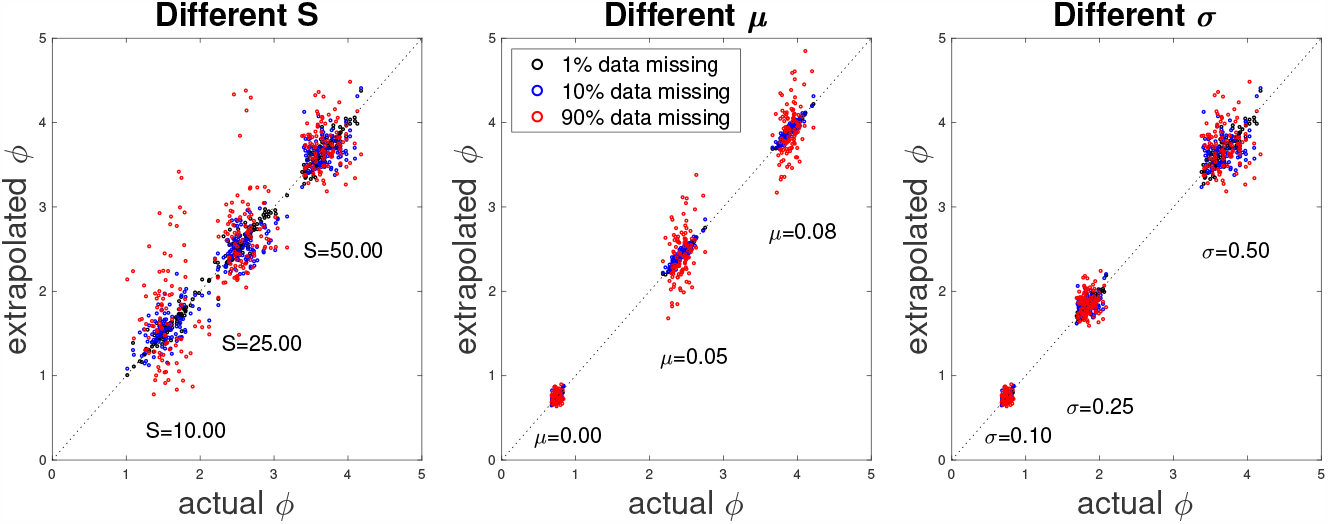
Estimating the collectivity parameter *ϕ* from incomplete data (in the absence of structure). We generate interaction matrices at random and compute their spectral radius (actual *ϕ*). For each we use a given fraction of their elements to draw the remaining fraction before computing the spectral radius (extrapolated *ϕ*). The clear pattern that we see here is that, as long as the size of the matrix is large enough, even a small fraction of interactions are enough to reliably estimate *ϕ*. This can be explained from the fact that the spectral radius of large random matrices only depend on basic summary statistics of elements.

**Figure S5:**
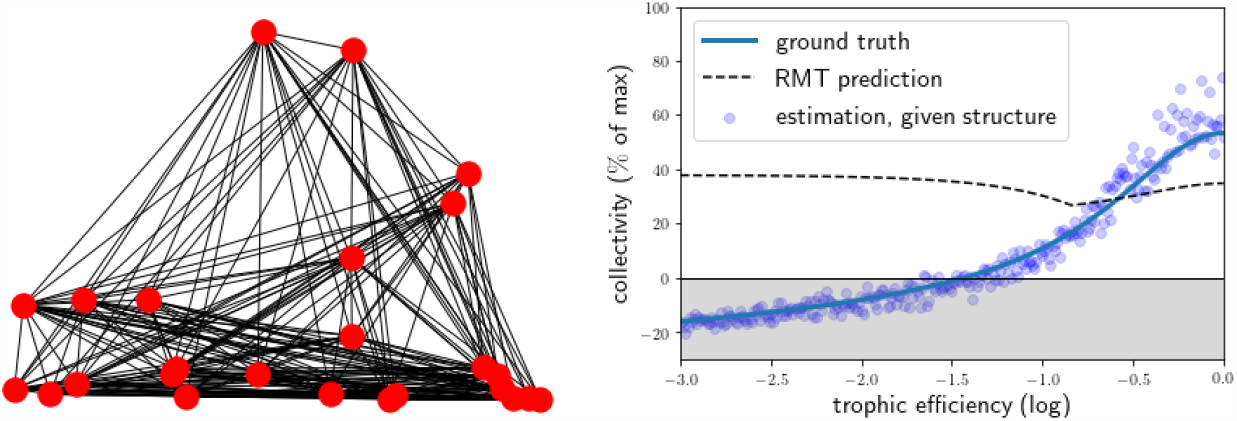
Effect of structure on the robustness of collectivity to noisy data. In cascade food web models, we modulate structure –from quasi triangular to perfectly anti-symmetrical– by varying trophic efficiency. Connectance and number of species are fixed, each point is a random realisation of the model, with different mean attack rate and trophic efficiency. As trophic efficiency grows, we go from a low degree of collectivity when systems are quasi-triangular (because upwards feedbacks are dampened) to highly collective when they become anti-symmetric. The fact that blue points lie along a thin band (near the reference values –the line– that we try to estimate) implies that collectivity is mostly driven by a combination of summary description of structure (the cascade topology and trophic efficiency) and interaction strength (statistics of attack rate).

This gives us an estimation of *A* from which we can deduce an estimation of *ϕ*. In Fig. S6 we generate many communities based or random competitive GLV models. We see on the left the result of the ansatz for one community. What is obvious here is that interactions are poorly predicted, with many false positives. The estimated spectral radius however is not far from the actual one. On the right we confirm that this was not just a lucky pick, collectivity in our examples is always well predicted. This is relates to our first point above, basic summary statistics of the interaction network can be sufficient if the community does not show particular structure. This is just a proof of concept however, showing that the tenuous signatures of interactions that may be present in correlation data can sometimes be enough to estimate collectivity. Making this point more precise and directly testable on data is a challenging research direction.

## S3 On the arbitrary bounds of communities

By ‘community’, we mean a set of interacting species, which could represent part or all of an ecosystem (Levins, 1974). It is most likely that the community does not exhaust the whole ecosystem in which it is embedded (in fact the latter would be hard to define). An ecologist might consider a community of plant species in a grassland, knowing of course that this grassland also contains insect species, fungi, bacteria, birds and larger herbivores that do have an influence on the plants. Those unobserved species (or abiotic compartments) will mediate the interactions between the species that we do consider explicitly, so that the interaction between two species is not an intrinsic characteristic of these populations, but is context-dependent. Thus, each interaction term is a property of the system, in a given state, rather than a fixed property of a given species pair.

The notion of “carrying capacity” –the abundance of a species if alone– must also be understood as context-dependent and not at all intrinsic to the focal species (Loreau, 2010). Carrying capacity changes as we change the scope of the community we are considering. If we only consider a single species, it must have a positive carrying capacity, otherwise it would be absent. But this carrying capacity might be an emergent feature of the myriad other species and abiotic compartment that we did not consider. As we increase the scope of the community under consideration, carrying capacity might become more intrinsic, but will most likely also decline. In the limit of all organisms on Earth, the notion itself ceases to make sense.

**Figure S6:**
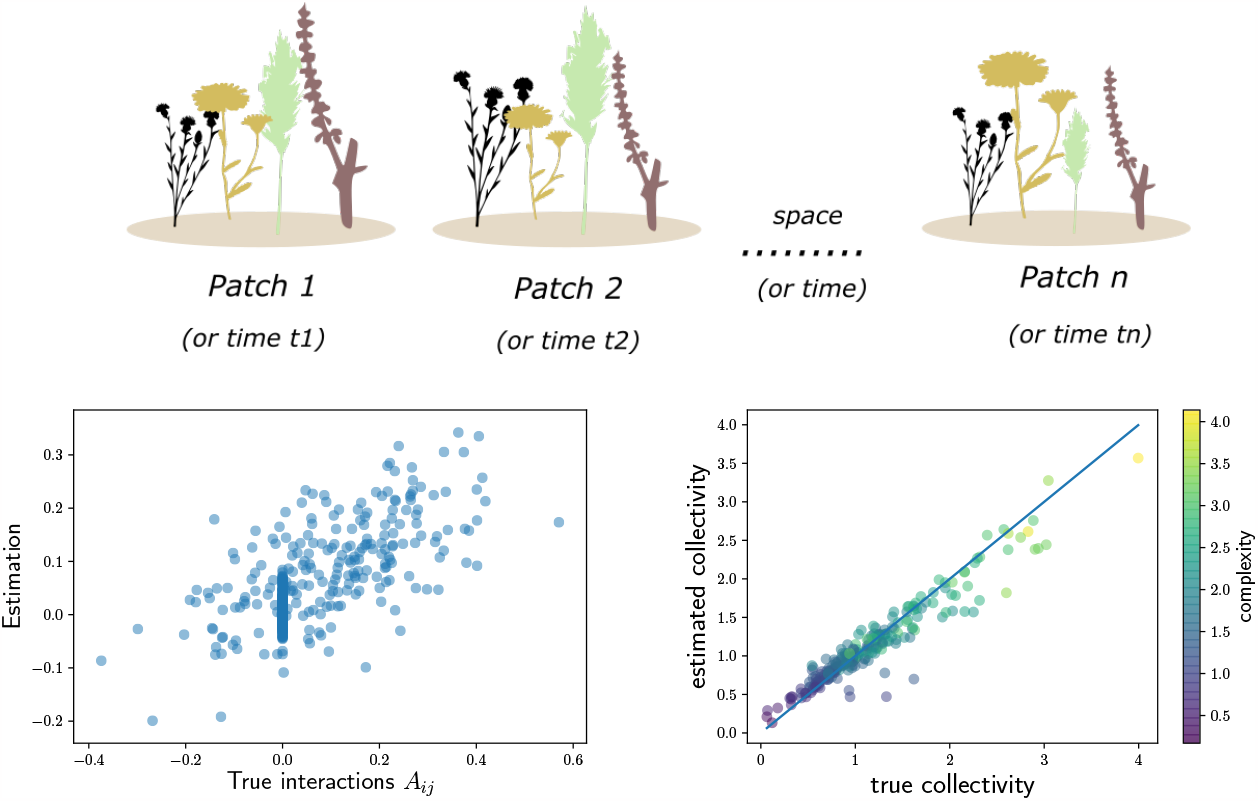
Estimating collectivity from species abundance correlations across patches. As a proof of concept for the reliable estimation of collectivity from empirical data, we generated many competitive communities, and for each assumed small variations in environmental conditions across patches. Using only the observation of the resulting species correlation matrix across patches we can guess species interactions (see ansatz in Eq. (27)) and thus collectivity. Left is an example of this guess, which we see is a very poor predictor on pair-wise interactions. Right: yet the estimation of collectivity is surprisingly good.

## S4 Indirect *vs* higher-order interactions

Higher-order interactions generalize pair-wise interactions to include more than two interacting partners at once. For example, a planktivorous fish might prey on a zooplankton species in the absence of another predatory fish, but not in its presence. This notion challenges the relevance of viewing communities as a graph, with nodes connected by directed edges (Battiston *et al*., 2021), since the existence and/or weight of an edge between two nodes is conditioned on the status of other nodes. We focus here on indirect interactions, and despite a very similar terminology, they are not equivalent to higher-order interactions. The latter occur when considering non linear extensions of GLV models, so that the coupling terms between dynamical variables depend on the product of several variables at once. Studying higher order interactions amounts to understanding the effects of such non-linearities. For us, this implies that the interaction matrix, as defined in Box 1 could be state-dependent, and not just dependent on the species composition (as is the case for GLV models). As is made clear in the formal derivation proposed in Box 1, this does not affect the definition of collective integration. From this perspective, our work may offer a novel direction of investigation: understand how higher order interactions affect the collectivity of steady sates.

## S5 Community assembly simulations

### S5.1 Model

The presentation of signatures of collective integration in Figures 2-5 are all based on the same set of model communities. This is a set of Generalized Lotka-Volterra (GLV) communities with random carrying capacities and interactions. The equations simulated are given by eq. 28

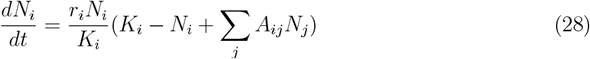

Where the (*K*_*i*_) are the species carrying capacities and (*r*_*i*_) their intrinsic growth rates. For simplicity we will choose *r*_*i*_*/K*_*i*_ ≡1, which amounts to enforcing an r-K tradeoff (but this is insubstantial to the points we make). We then consider a gradient of interaction strength, with 50 different values of average interaction strength along the gradient, each with 100 random communities, making up 5000 communities in total. Each community starts with 50 species (stronger interaction leads some species to extinction in the equilibrium state), and we set 80% of the interactions to zero, so that we have a sparse interaction matrix. For defining the interaction matrix, we follow (Barbier *et al*., 2018) and define the three parameters of random interactions as *μ* =*−y*; *σ* = *y*; *γ* =*−*1, where *y* is the above-mentioned gradient, with values ranging between 0.02 and 1.0, in uniform intervals.

This translates to the following relations. The carrying capacities *K*_*i*_ are taken from a random uniform distribution centered on 1, with a standard deviation of 0.2 (within the span of 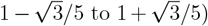. The interactions matrix *A*_*ij*_ is taken from a Gaussian distribution, except for the diagonal which is set to *−*1. For the off diagonal values, we constrain the values to keep three conditions: i) asymmetrical interactions, so that *A*_*ij*_ =*−A*_*ji*_; ii) we use a sparsity of *ρ* = 0.8, i.e. 80% of of the interactions values are set to 0; iii) the average and standard deviation of the interaction values (including the values set to 0, but not the diagonal values) are *−y·S*, 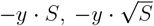, respectively (*S* = 50 is the number of species in the community).

These constraints are achieved by building a matrix in the following order: 1) take a matrix *A*_1_ of size *S* from a random Gaussian distribution; 2) Get a new matrix by subtracting the original matrix from its transpose: 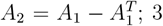) normalize the matrix by its standard deviation: *A*_3_ = *A*_2_*/std*(*A*_2_); 4) use the normalized matrix to get a certain average and standard deviation: 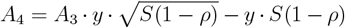; 5) Choose at random a fraction of *ρ* of the upper triangualar part of the matrix, and set it to 0, as well as its lower triangualar counterpart.

This gives us asymmetrical interactions between species (*γ* =*−*1), and interactions that are increasingly negative (due to *μ*) and at the same time more varied (due to *σ*). Using this definitions, along the gradient we find communities with a collectivity parameter *ϕ* ranging between 0 and 2, allowing us to showcase different aspects of low and high collective integration in communities.

For each of the 5000 communities, we use the carrying capacity vector *K*_*i*_ and interaction matrix *A*_*ij*_ to assemble a community. We run a simulation until reaching an equilibrium (as detailed in the next subsection) with the starting conditions that all 50 species abundances equal 1. Depending on the location along the gradient, all species survive for low values of *y*, or at least 30 species survive at high *y* values. For any given community, we consider only the extant species for calculations and presentation (e.g. *ϕ* is calculated for the *A*_*ij*_ matrix where all rows and columns of extinct species are taken out).

### S5.2 Simulations

The simulations of the ordinary differential equations (ODEs) given by eqs. 1 were preformed using the ode45 solver of Matlab (R2019a). The computer code used for the simulations is found at the open-access repository: https://doi.org/10.5281.

To reach an equilibrium we start with a simulation time span of *T* = *T*_0_, and iteratively run a simulation for longer time spans of *T* (doubling *T*) until the equilibrium condition defined below is reached, or until we reach a maximal simulation time *T*_*max*_. The equilibrium condition is that at the last quarter of the simulation the maximal change in species abundance is below

the threshold *z*, scaled by the time span *T*, max(abs(*N*_*i*_(*T*)*−N*_*i*_(0.75*T*))) *<* 2*zT*. We choose the numerical parameters *T*_0_ = 10^2^, *T*_*max*_ = 10^4^, *z* = 10^*−*6^, ensuring that the abundances no longer change in any noticeable way once reaching the threshold.

## S6 Presentation of signatures

We describe here in detail the calculations and simulations used for the presentation of signatures of collective integration in Fig. 2-4, as well as an additional signature not described in the main text: the increase in effective connectance (see Fig. S7).

### S6.1 Effective connectance

If a community is collectively integrated, substantial net interactions should connect any two species, even if these species do not directly interact. In other words, the effective connectance of the community should be much larger than that of the direct interaction network. The signature of effective connectance is defined using a ratio between two Shannon diversity index, applied on two matrices that describe the interactions within the community. We use the set of communities as described in the previous subsections, and make use of two matrices: the direct interactions matrix *A*_*ij*_, and the net interactions matrix, *V*_*ij*_ = (1*−A*_*ij*_)^*−*1^. To calculate the effective connectance we use the ratio between the Shannon diversity of these matrices, calculated as follows.

For each matrix we calculate a diversity of interactions by: 1) taking all interactions except self-interactions (i.e. diagonal values), giving us *S*(*S−*1) values; 2) computing the prevalence of each unique value as a fraction *ρ* = *x/S*(*S−*1), where *x* is the number of occurrences of each value; 3) calculating the Shannon diversity on these values as *H* =*−ρ ·log*(*ρ*). We then define effective connectance as the ratio of the two indices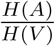. Note that for our definition of unique values, we round all values by 10^*−*^3 (e.g. all values between 0 and 10^*−*^3 are considered the same, all values between 10^*−*^3 and 2*10^*−*^3 are considered the same, and so on), as otherwise all random values will be effectively unique and the definition of Shannon diversity would be rendered meaningless. The result of this calculation is used for the y-axis measure of Fig. S7a. For Fig. S7b-d we show the histograms of off-diagonal values for both *A*_*i*_*j* and *V*_*i*_*j*, for three specific communities along the *y* gradient.

**Figure S7:**
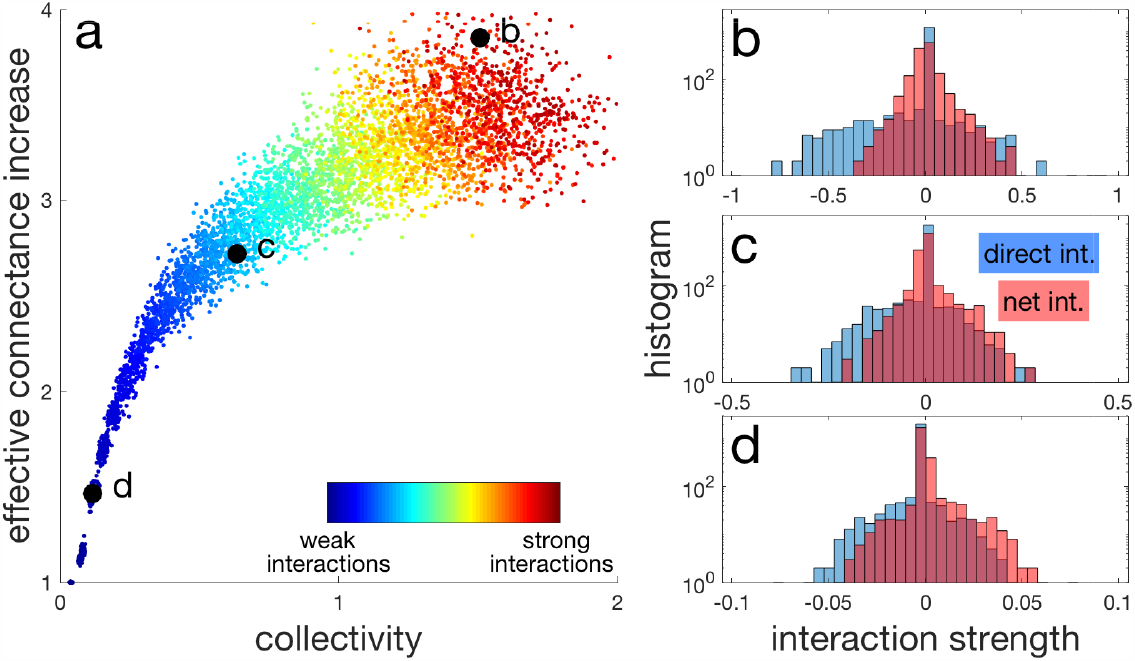
Growth in the effective number of net interactions per species, relative to the number of direct ones, as collectivity increases. Left panel shows the effective connectance – the ratio between the Shannon diversity index of the direct interactions and net interactions, as defined in the main text. Black circles highlight several communities that are considered in the right panels. The right panels show the histograms of direct and net interactions, overlayed, for these communities. Note the logarithmic scale on the y-axis and the changes across panels of the x-axis.

### S6.2 Perturbation depth

The signature of perturbation depth is tied to the response of the community to an external perturbation, and we therefore calculate it using a dynamical simulation (as described in the previous subsections). For a given community at its equilibrium abundance 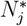, we consider *S* different scenarios, in each of which we eliminate one species from the community, and observe the effect on the remaining *S −* 1 species. We simulate the community dynamics until it reaches a new equilibrium, and then measure the change in abundance for all *S −* 1 species, 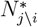. For this community, we also note the link distance *d*(*i, j*) between two species *i* and *j*, where a distance of 1 is for directly interacting species, distance 2 is for species that do not directly interact but that both interact directly with the same third species, and so forth. We these ingredients we calculate the perturbation depth as:

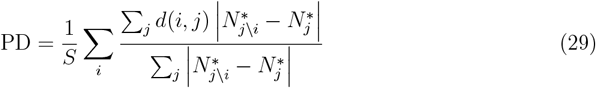

Note that in these summations over *j* we go through all species except for the removed species, whereas the summation over *i* is essentially over different removal experiments. The result of this calculation is used for the y-axis measure of Fig. 2a. For Fig. 2b we calculate the relative impact of the perturbation of three groups of species, the ones directly interacting with the removed species (red), the one that interact with a directly interacting species (green), and the rest of the species in the community that are at a distance of 3 or more (blue). This relative impact is the ratio between the change in abundance within the group, and the change in abundance for all the species in the community (except the removed one), and all of this is averaged over the different removal experiments.

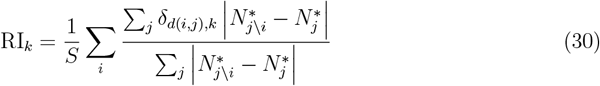

**Figure S8:**
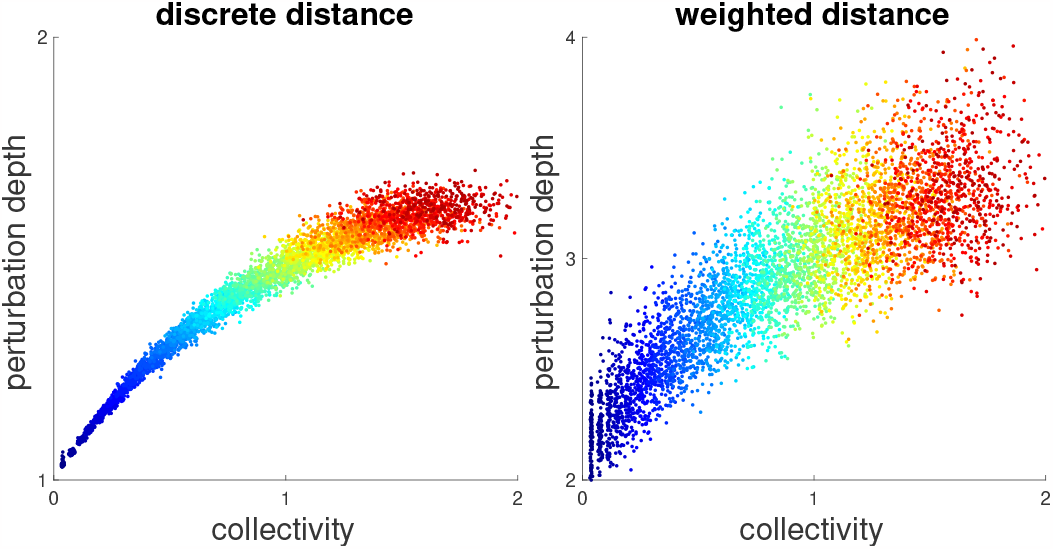
Estimating perturbation depth using two different methods of defining distance in a network. Both panels show perturbation depth as a function of collectivity *ϕ* (as defined in the main text), with the left panel using the same definition of distance as in the main text, while in the right panel we use a weighted distance definition, that can be used in non-sparse networks as well.

#### S6.2.1 Generalizing perturbation depth beyond sparse networks

In the main text we defined perturbation depth in a way that requires the interaction network to be sparse, as this allowed for a simple an straight-forward definition and presentation of the signature. We can, however, also define perturbation depth using the quantitative values of the interaction in another way, so that it can be defined on all interaction matrices. In the right panel of Fig. S8, we show results for this method (compared to the original results on the left). In this alternative method, network distance is the shortest path between two nodes following edges of the network, where the length of an edge reflects the (two-way) interaction strength between the species that it connects. In this way we can define a notion of perturbation depth that can be applied beyond sparse networks. The length to every edge of the network is chosen (somewhat arbitrarily) as

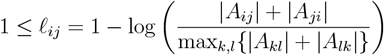

with the convention that for two species that do not interact, ℓ_*ij*_ =*∞*. We then numerically (using the dijkstra algorithm) find the shortest path between nodes according to the above edges’ length. We finally use this shortest path as our distance measure for calculating perturbation depth.

### S6.3 Temporal unpredictability

The signature of temporal unpredictability relates to the difference between short-term and long term response, and as such we use dynamical simulations both up to equilibrium, and also within a short timescale.

We look at how the community responds to perturbations by changing the carrying capacity of each species at random: 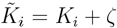 where *ζ*is chosen at random from a uniform distribution centered around 0, ranging from *−*0.25 to 0.25. We then run a simulation twice, first over a short time span Δ*T* = 0.5, and then until the system reaches a new equilibrium 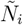. From the first simulation, for each species separately, we extract 10 time points (regularly spaced) within Δ*T*, and fit the points to a linear function. We use the slope *α*_*i*_ from the fit to extrapolate what the long term response is expected to be: *R*_*S*_ = *α*_*i ·*_*K*_*i*_*/*(*r*_*i*_*N*_*i*_). This is compared to the actual long term response 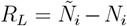, giving us our signature of temporal unpredictability, as defined by eq. 31.

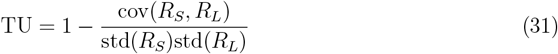

The result of this calculation is used for the y-axis measure of Fig. 3a. For Fig. 3b-c we also demonstrate the dynamics of two communities along the *y* gradient. Fig. 3b-c we show the actual long tern dynamics (solid lines), compared with the predicted long term dynamics, extrapolated from the short term (dotted line). The extrapolated curves shown are: Δ*Ñ*_*i*_(*t*) = (1 *−* exp(*−r*_*i*_*N*_*i*_*/K*_*i*_*t*)) *· α*_*i*_*K*_*i*_*/*(*r*_*i*_*N*_*i*_).

### S6.4 Biotic contribution to the realized niche

Biotic contribution (BC) measures how much the abundances are determined by the community as a whole, rather than by the abiotic conditions alone. We measure it using the relative yield of species *η*_*i*_ = *N*_*i*_*/K*_*i*_, where we use the equilibrium abundances *N*_*i*_. This gives us the signature, used for the y-axis of Fig. 4a, as

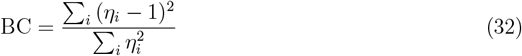

For Fig. 4b-d, we use for the x-axis the carrying capacities *K*_*i*_ (blue), and the term 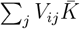 which represents the effects of the community on the species (red), where we sum over the matrix of net interactions *V* = (𝕀 *− A*)^*−*1^, and 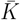 is the mean carrying capacity.

## Footnotes

1 The Jacobian matrix that controls near-equilibrium dynamics reads *J* = *D*_*N*_ (*−*𝕀+ *A*), where *D*_*N*_ is a diagonal matrix whose entries are 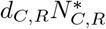, and 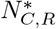 are the species equilibrium values. Stability requires that eigenvalues of *J* have negative real part. Here we only know that eigenvalues of *−*𝕀+ *A* have negative real part, but we can always adjust the positive factor *D*_*N*_, by modifying *r* and *m*, so that it will not change the sign of eigenvalues.

2 for instance, *ϕ* is equal to the upper bound in a competitive network where competition strength is uniform across species (See also Appendix S1).

3 The Jacobian matrix can be written as *J* = *D*_*N*_ (*−*𝕀+ *A*), the product of a *positive* diagonal matrix *D*_*N*_ that contains species abundances at steady state and their intraspecific interaction strength, times our non-dimensional interaction matrix. In general *J* and *−*𝕀 + *A* will not have the same eigenvalues, but because *D*_*N*_ is diagonal and positive, it is unlikely (although not impossible) that some eigenvalues’ real parts become positive when going from *−*I + *A* to *J* (Gibbs *et al*., 2018).

4 Those cases are when *E∝I* (no structure in the environmental variations) and *A* is positive definite (which implies symmetric interactions).

## Notes

https://doi.org/10.5281/zenodo.7537451

### Competing Interest Statement

The authors have declared no competing interest.

### Summary of Updates

This version has been updated substantially to improve the framing, and explain why our theory can be used to quantify the dynamical role of interaction structure. We provide inn this new version examples of structured community models, such as food-webs.

